# A Compact Matrix Representation for Hydrocarbon and Its Applications on Monoterpene Carbocation Rearrangement Elementary Steps

**DOI:** 10.1101/2025.08.23.671925

**Authors:** Tianbo Tse

## Abstract

This work proposes a compact matrix-based representation for hydrocarbon carbocations ([C_*x*_H_*y*_]^*m*+^). In addition, its application is explored for the elementary steps of non-stereoisomeric [C_10_H_17_]^+^ rearrangement.

## 1 Introduction

Rapid development of Artificial Intelligence for Science (AI4S) has led to the emergence of numerous molecular descriptors that enable efficient machine learning applications in chemistry[1]. Among these, the adjacency matrix remains one of the most classical representations of molecular structures. We proposed a compact and specific optimized adjacency matrix for hydrocarbon backbones.

## 2 Adjacency Matrix Representation Review

### 2.1 Basics of Adjacency Matrix

The classic graph-based method for molecular representation is adjacency matrix.

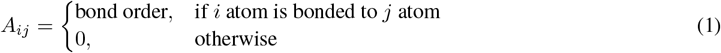

We will use ethane cation as an example(see Figure 1), we labeled atoms from 1 to 7. It is obvious that the largest number is the number of atoms in the molecule. Since bonding is commutual, the adjacency matrix should be symmetric.

**Figure 1.**
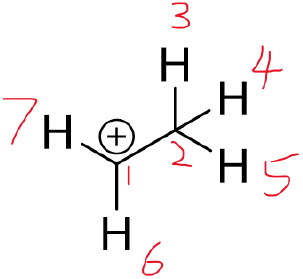
Ethane Cation with Labeling.

### 2.2 Only Heavy Atom Consideration

In this case, only carbon atoms will be considered. That will cause a compact 2 x 2 matrix:

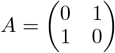

However, critical information is lost in this representation. Ethane will share the same representation as the ethane cation. In other words, the number of hydrogen and charge information is not explicitly shown in this representation. Hydrogen/charges are critical in a mechanism proposal or in other theoretical chemical analysis.

### 2.3 Full Atom Consideration

To overcome the information lost by only heavy atom consideration, a full atom consideration is a natural extension:

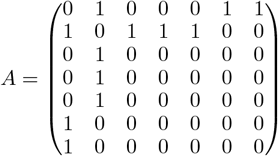

The full atom consideration solved the hydrogen missing problem, but charge information is still missing. Also,this matrix is very spouse. The density of information is not ideal.

### 2.4 Intrinsic Weaknesses about Adjacency Matrix Representation

- The diagonal terms are always 0. Trace should be an important parameter for a matrix.
- Methyl/Ethyl groups can lead same row/column of a matrix, leading a 0 determinant.
- Most of dimensions are contributed by hydrogens, leading a very spouse matrix.
- The charge is not shown in the matrix.

## 3 A Compact Representation for Hydrocarbon (Cation)

### 3.1 Construction Rules

To overcome intrinsic weaknesses of adjacency matrix representation. We proposed simple rules for a compact, information lossless representation.

- Label all carbons from 1 to j, j is the number of carbon items.
- *M*_(_*ii*) indicates the number of hydrogens attached on the i th carbon

Under such representation, the positive charge can be obtained by calculating the sum of a row/column because carbons with formal charge will only have 3 bonds, but saturated carbon atoms have 4 bonds.

### 3.2 Example – Ethane Cation

Our starting point is only heavy atom consideration, but all hydrogens can be considered together. There are 2H attached to C1 and 3H attached to C2, following the above rules, we can get:

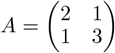

For charge information, we can calculate the sum of specific row/column, it does not matter we choose row or column because this matrix will be symmetrical. Based on calculation, carbon 1 carries the positive charge.

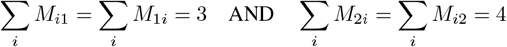

### 3.3 Shallow Intrinsic Properties of Such Representation

- Trace of the matrix is the number of hydrogens in the molecule.
- Non-Zero determinant can be obtained, that could be a more compact representation for a system(**Degeneracy should be considered!**)
- Dimension of the matrix is always the number of carbon.

### 3.4 Permutation Group Structure of such Representation

The way of labeling should not change any information about the molecule. For example, the following matrix should also represent the ethane cation.There should be *N* ! permutation representation for a N carbon containing hydrocarbon molecule and all permutation group theory could be applied in such representation. One practical invariant for a matrix with line/row permutation is the determinant. Another useful invariant is trace of any power of the matrix. These can be kind of unique for

### 3.5 Physical Interpretation for Such Representation – Benzene

M matrix is very similar to the tight binding model or Hückel method Hamiltonian of benzene. The main difference is on off-diagonal terms, in our representation they are positive, but in tight-binding model, they are negative.

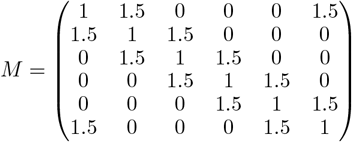

We can calculate the eigenvalue(energy spectrum) and eigenvector(molecular orbital coefficient) of M and H matrix, we will get almost same results but opposite correspondence between energy and orbital coefficient. Such opposite correspondence is caused by positive off diagonal terms. We can interpreted this representation as a modified tight binding model.

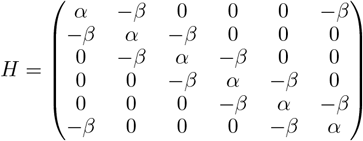

## 4 Applications on Monoterpene Carbocation Rearrangement Elementary Steps

Terpene carboncation backbone is an ideal platform for our representation application due to the terpene biosynthesis mechanism. We applied our representation to four types of elementary steps of monoterpene carbocation rearrangement considered by Jacobson and coworkers [2].

### 4.1 Starting Material and Its Matrix Representation

The starting material is given by Figure 3. It has clear chemical structure(no complex topology structure) and it is a good platform to test several elementary steps for a carbocation backbone.

**Figure 2.**
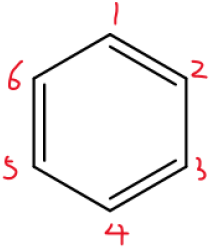
Benzene with Labeling.

**Figure 3.**
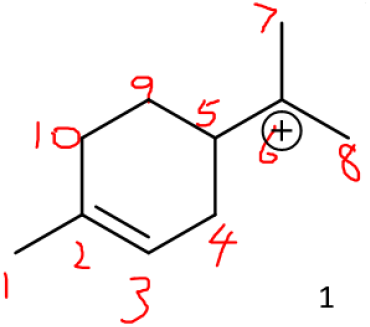
Starting Material with Labeling.

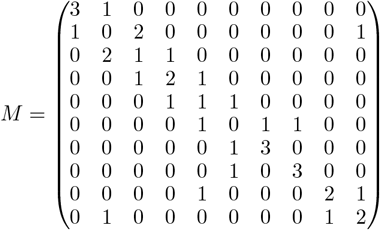

### 4.2 Intramolecular Alkylation of Double Bonds

The key part for intramolecular alkylation of double bonds is to identify the carbocation and double bonds. To identify carbocation, we can sum over all elements in one column or one row. The 4 indicates a saturated carbon, 3 indicates a carbocation. C6 is the carboncation in the starting material. Identifying double bond(s) is easier, the off diagonal 2 elements are targets. We can easily find that the double bond is between C2 and C3.

**Figure 4.**
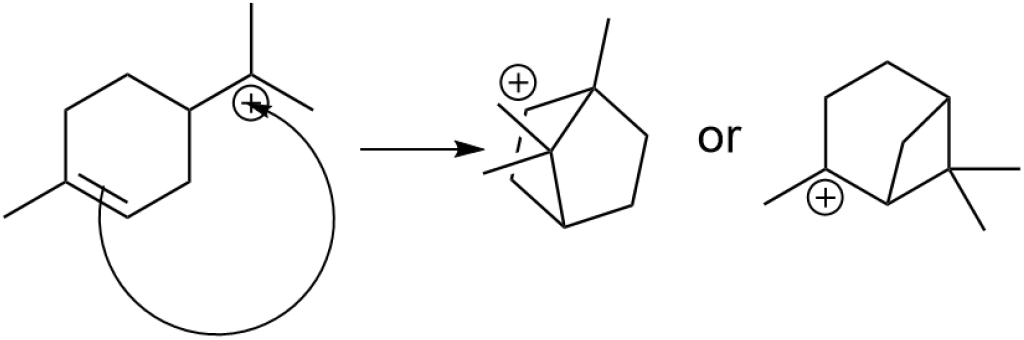
Intramolecular Alkylation of Double Bonds.

The transformation is straightforward:

- Reduce off diagonal terms from 2 to 1 (double bond to single bond)
- Change *M*_*ij*_ and *M*_*ji*_ from 0 to 1 (*i* is the place of double bond, *j* is place of carbocation)
- There should be at least 2 products since double bonds are commutual

### 4.3 1,2-Alkyl Shift

The workflow for 1,2 alkyl shift is a little bit different; we have to identify the “distance 2 carbon,,“ which means the carbon that is 2 bonds away from thecarbon cationon.

**Figure 5.**
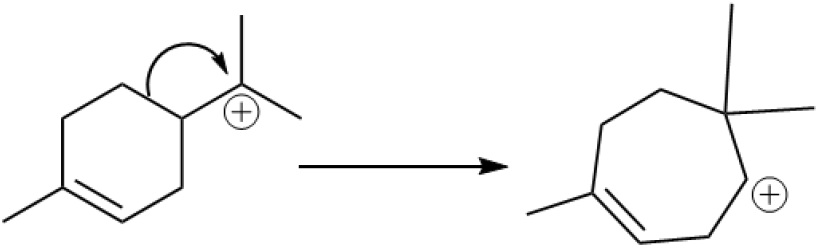
1,2 Alkyl Shift.

1. Identify distance 2 carbons and distance 1 carbons
2. Break the bond between distance 2 carbon and distance 1 carbon
3. Form a bond between carbon 2 and carbocation
4. Theoretically, there are at most 4 different products of 1,2 alkyl shift
5. Methyl shift is a specific 1,2 alkyl shift

### 4.4 Hydride Shift

Only 1,2 hydride shifts are considered here because other hydride shifts can be described as series of 1,2 hydride shift.

**Figure 6.**
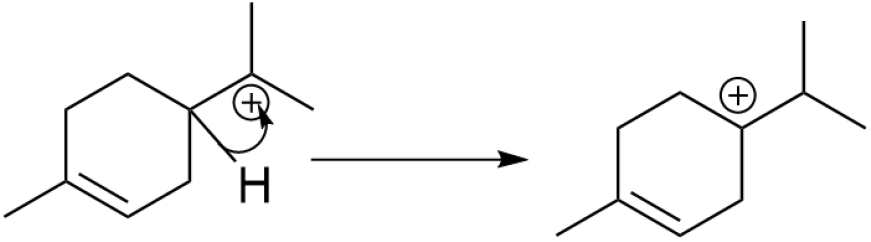
Hydride Shift.

1. Identify the distance 1 carbons, make diagonal term for that carbon minus 1 (lose 1 H)
2. Add 1 on diagonal term for carbocation, make it neutral.

### 4.5 Proton Transfer

The product of proton transfer is versatile, theoretical there are 3 possible distance 1 carbon to be deprotonated and 2 possible carbon can gain that proton.

1. Identify distance 1 carbon and remove 1 H from distance 1 carbon
2. Make distance 1 carbon and carboncation double bond
3. Make original double bond to single bond
4. Chose 1 of 2 double bond carbon to gain the proton

**Figure 7.**
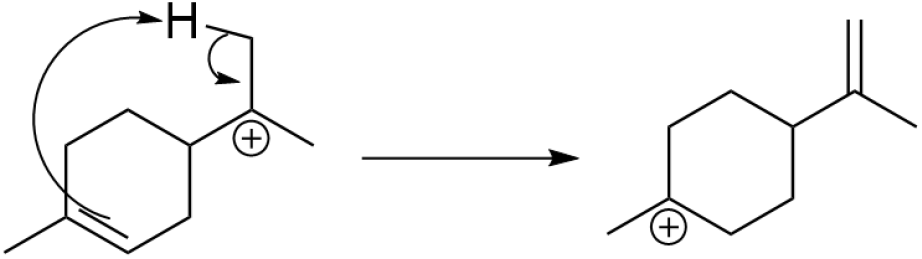
Proton Transfer.

## 5 Limitation

The limitations listed here are mostly inherited from adjacency matrix representation. Unfortunately, this work has not solved these limitations.

– The stereochemistry/anion/radical is not considered.
– This representation is specified for hydrocarbons, and common elements like N are not supported.
– The operation is based on specific elements operation, not on the matrix product.

## 6 Conclusion

This work proposed a compact matrix based representation for hydrocarbon ([C_*x*_H_*y*_]^*m*+^). Its application for elementary steps of non-stereoisomer [C_10_H_17_]^+^ rearrangement is explored. Limitations about this representation are pointed out. We believe that such representation can contribute on AI for chemistry because its strong physical meaning.

## Special Acknowledgment

The author thank Dr. Rachel R. Schendel (Department of Animal & Food Sciences, University of Kentucky) for her kind support during author’s academic study at University of Kentucky.

## Notes

### Competing Interest Statement

The authors have declared no competing interest.

